# Renal ischemia-reperfusion injury attenuated by exosomes extracted from splenic ischemic preconditioning

**DOI:** 10.1101/2022.06.25.497584

**Authors:** Liu Hongtao, Shen Ye

## Abstract

**Objective:** To investigate the protective effects of the exosomes extracted from splenic ischemic preconditioning (sIPC) on renal ischemia-reperfusion (IR) injury.

**Materials and methods:** Splenic ischemic preconditioning(sIPC)was conducted on mice in vivo 24 hours before the start of renal ischemia-reperfusion (IR) injury experiment, and serum exosomes derived from sIPC mice were infused into the mice model of renal ischemia-reperfusion injury. The kidney tissue and serum were collected 24 hours later. The morphological changes and inflammation in ischemia-reperfusion kidneys were determined by hematoxylin-eosin (HE) staining.Then the apoptosis of kidney tissue sections were detected by TUNEL staining, Ki-67 immunohistochemical staining was used to assess the proliferation.In addition, the levels of pro-inflammatory cytokines including TNF-α, IL-1β and SCr in serum were measured by ELISA.

In vitro, we extracted exosomes from mouse spleen fibroblasts pretreated with hypoxia and reoxygenation (H/R) and administered them to mouse renal epithelial cells.Furthermore, for the hypoxia-reoxygenation model of renal epithelial cells, TUNEL and flow cytometry were used to evalutaed cell apoptosis;Then ELISA was used to measure the levels of TNF-α and IL-1β in the cell supernatant, Bax and Bcl-2 were measured by Western Blotting.

**Results:** HE staining showed that the renal injury caused by ischemia-reperfusion attenuated after sIPC. TUNEL staining showed that renal tissue apoptosis was greatly reduced after sIPC or injection of exosomes extracted from splenic fibroblast hypoxia-reoxygenation model. Ki-67 staining showed that the positive rates of IRI+sIPC group, IRI+mSF(H/R)-exo group, IRI+mSF(H/R+PBS)-exo group were close, higher than IRI group but lower than sham group. ELISA test of kidney tissue showed that the serum creatinine, TNF-α and IL-1β induced by IRI decreased with sIPC and addition of the above-mentioned exosomes.In vitro, the exosomes extracted from the hypoxia-reoxygenation model of splenic fibroblasts had the same protective effect on hypoxia-reoxygenated mouse renal epithelial cells model, and this protective effect disappears after the addition of exosome inhibitors.TUNEL and flow cytometry showed that the exosomes reduced the apoptosis. The ELISA test results showed that the levels of TNF-α and IL-1β in the H/R group increased significantly, but decreased due to the splenic fibroblast exosomes treated with starvation.While the exosome inhibitors inhibited the effects of exosomes.Western blot results showed that the Bax expression level of the H/R group increased, and the Bcl-2 decreased.While the starvation-treated splenic fibroblast exosomes decreased the Bax level and increased the Bcl-2 level.

**Conclusions:** The exosomes extracted from splenic ischemic preconditioning exerted a protective capacity to attenuate renal IR injury.

## Introduction

Renal ischemia reperfusion triggers a series of oxidative and inflammatory reactions, which is the main cause of chronic and acute renal failure and the reason which causes poor graft survival rate after renal allograft transplantation. Remote ischemic preconditioning refers to interspersing 3-4 remote local ischemia treatments before the continuous ischemia of the heart, brain, kidney and other tissues to protect important organs and tissues from continuous ischemia-reperfusion-induced injury^[1]^. As the body’s largest immune organ, the spleen is the center of the body’s cellular and humoral immunity. It plays an important role in the IRI of myocardium, lung tissue, and kidney ^[2-4]^. Interestingly, the renal function of rats that underwent splenectomy before renal ischemia-reperfusion was better^[5]^.Our previous study also have shown that splenic ischemic preconditioning reduces renal ischemia-reperfusion injury^[6]^. Exosomes are vesicle structures released by various cells under physiological conditions or disease states ^[7]^. The nucleic acids^[8]^, proteins^[9]^ and lipids in the exosomes can serve as media for information dissemination, thereby affecting the function of recipient cells. Pan et al found that the miR-21 derived from exosomes in the serum of mice with remote ischemic preconditioning can alleviate the septic acute kidney injury^[10].^

In this work, we hypothesized that splenic ischemic preconditioning can act on the kidneys through exosomes to further reduce the renal IR injury. We applied the exosomes extracted from the hypoxia/reoxygenation model of splenic fibroblasts and exosomal inhibitors to the mice model of renal ischemia-reperfusion injury and the mice renal epithelial cell model of hypoxia/reoxygenation, respectively, and assessed the effect of exosomes on renal ischemia-reperfusion injury. The possible mechanism was also explored. Our research may provide a potential therapeutic modality for renal ischemia-reperfusion injury.

## Materials and Methods

### Animals

All 6 to 8 week-old male C57BL/6J mice were provided by the Changzhou Cavens Laboratory Animal Co., Ltd to establish animal models.The animal study protocol was in accordance with the Guide on Care and Use of Laboratory Animals published by the National Institute of Health and approved by the Ethic Committee of the People’s Hospital of Subei. All mice were administered with free access to food, and water in a clean environment at room temperature of 20-23°C and fasted for 12 hours before the operation.

### Surgery procedure

The mice were randomly divided into seven groups (n=6 for each group): the sham operation group (sham), the renal ischemia-reperfusion injury group (IRI) and renal ischemia-reperfusion injury after the splenic ischemic preconditioning and reperfusion group (IRI+sIPC),exosomes derived from splenic fibroblasts and renal ischemia-reperfusion injury group (IRI+mSF-exo),exosomes derived from mouse splenic fibroblasts hypoxia/reoxygenation model and renal ischemia-reperfusion injury group (IRI+mSF(H/R)-exo),exosomes derived from splenic fibroblasts hypoxia/reoxygenation model with inhibitor and renal ischemia-reperfusion injury group (IRI+mSF(H/R+GW4869)-exo),exosomes derived from splenic fibroblasts hypoxia/reoxygenation model with phosphate balanced solution and renal ischemia-reperfusion injury group (IRI+mSF(H/R+PBS)-exo).

For all the groups, the animals were anesthetized with chloral hydrate intraperitoneally in a dose of 125 mg/kg. In the sham group, the abdominal cavity was opened and the renal pedicle was separated, but not clamped. After 20 minutes, the abdominal cavity was closed by layered sutures.For the IRI group, the left renal pedicle was clamped with non-traumatic vascular clips for 20 minutes.It can be seen that the kidney changes from bright red to purple-black, indicating clamped successfully; then, remove the clips and restore blood perfusion, it can be seen that the kidney quickly return to their original color. After the operation, the abdominal cavity was closed. For the IRI+sIPC group, three cycles of splenic ischemic preconditioning were firstly performed by using non-traumatic vascular clips before the renal ischemia: the blood vessels in the splenic pedicle were dissected and, clamped to ensure the blockage of blood flow. Each cycle of the splenic ischemic preconditioning was 5 min of ischemia followed by 5 min of reperfusion. Following three cycles of splenic ischemic preconditioning, the renal ischemia was performed for 20 minutes as above. For the IRI+mSF-exo group, 150μL of exosomes which were extracted from normal-growing mouse splenic fibroblasts (mSF) was injected into the tail vein and then the renal ischemia was performed for 20 minutes.For the IRI+mSF(H/R)-exo group, the exosomes derived from mouse splenic fibroblasts hypoxia/reoxygenation model were extracted and injected.For the IRI+mSF(H/R+GW4869)-exo group, GW4869 which was an exosome inhibitor with a final concentration of 10μM was added to the exosomes extracted from the mSF(H/R) cell model.For the IRI+mSF(H/R+PBS)-exo group, GW4869 was replaced with PBS. For all the groups, the mice were reared in the environment of 24∼29 °C for 24h, supplemented with water and food in individual cage after the operation. All the mice underwent another anesthesia with the anesthetic protocol above after 24 h.The blood samples were collected from a puncture to the inferior vena cava and the kidneys were harvested for further experiment. Subsequently, a lethal dose of anesthetic was used for euthanasia of the mice.

### Cell model construction

We first established mouse splenic fibroblasts (mSF) hypoxia/reoxygenation (H/R) cell model as follows:For mSF that grow to 80% confluence, replace the culture medium with a sugar-free and serum-free medium, spread the sterilized liquid paraffin on the culture medium (0.24 mL/cm2), and then incubate at 37°C for 4 hours. Aspirate the paraffin on the surface, discard the medium, wash twice with PBS, add low-sugar 15% serum medium and continue to culture for 24 hours. Further, we established mouse renal epithelial cell hypoxia/reoxygenation (H/R) model using the same method to construct the mSF H/R cell model.

### Extraction of the exosomes

The cell supernatant was collected and mixed with 0.5 times volume of exosomal extraction reagent;The mixture was fully mixed and then incubated overnight in the 4 ° C refrigerator.The supernatant was abandoned after being centrifuged at 4 ° C for 1 hour, and the bottom was resuspended in an appropriate amount of PBS, which is an exosstally suspension. The extracted exosomes were observed by electron microscope.

### Histological analysis

The kidney tissues were fixed in 10% formalin, embedded in paraffin and sliced into 3µm sections. To analyze the histopathological changes, the sections were stained with hematoxylin-eosin. After deparaffinization with xylene and hydration with agraded ethanol series, the sections were soaked in hematoxylin for 10 min and washed with cold running water. Following incubation with 0.7% hydrochloric acid and ethanol for a few seconds, the sections were washed and stained with eosin dye solution.Then, all sections were dehydrated in ascending alcohol solutions (50%,70%,80%,twice with 95%,and twice with 100%) and xylene.

Further, the ki67 immunohistochemistry was used to reflect cell proliferation. The sections were deparaffinized and subjected to antigen retrieval with citrate buffer at pH 6. Then 3% hydrogen peroxide was used to block endogenous peroxidase.Following blocking with 3%BSA at room temperature for 25 min, then the samples were stained with anti-ki67 (1:1000) antibodies at 4°C overnight. Then the sections were incubated with goat anti-mouse secondary antibodies for conjugating with horseradish peroxidase (HRP) for 50min.3,3’-Diaminobenzidine(DAB) was used to detect the immunocomplexes, whereas hematoxylin was used for nuclear counterstaining.

The sections were examined by light microscopy(Nikon Corp.,Tokyo, Japan). Images were analyzed using Image J (Rawak Software Inc., Stuttgart, Germany). Three fields were selected in each slice randomly.

### Measurement of apoptosis and damage

#### Detection of kidney tissue damage using TUNEL assay

After baking in the oven at 60 °C for 60 min, the sections which were cut from the paraffin-embedded tissue were dewaxed with xylene twice for 5 min, hydrated using an ethanol gradient (twice with 100% for 3 min, then 95% for 3 min and 75% for 3 min), washed in PBS for 5 min and then incubated with proteinase K(20ug/ml)at room temperature for 15 min to remove tissue protein.Later, the sections were washed with PBS containing 2% hydrogen peroxide for 5 min at room temperature.Further, the TUNEL assay kit (KeyGEN BioTECH, Nanjing, China) containing TdT was used according to the manufacturer’s protocol.Finally, apoptotic cells in the sections were detected using a microscope (Nikon Corp.,Tokyo, Japan).

#### Detection of cell apoptosis using TUNEL assay

The mouse renal epithelial cells were fixed in 4% formaldehyde solution for 30 minutes at room temperature and then incubated with PBS containing 0.3% Triton X-100 at room temperature for 5 minutes.The TUNEL assay kit (KeyGEN BioTECH, Nanjing, China) containing TdT was used according to the manufacturer’s protocol. Subsequent to washing with PBS, the cells were counterstained with dAPI. Apoptotic cells in the sections were detected using a microscope (Olympus, Tokyo, Japan)

#### Detection of cell apoptosis using flow cytometry

The apoptosis of cells was examined using an Annexin V-FITC/PI apoptosis kit (Solarbio, Beijing, China) according to the manufacturer’s protocol. Briefly, the cells were collected and washed in pre-chilled phosphate-buffered saline (PBS) after digested with trypsin without Ethylenediaminetetraacetic acid (EDTA). Cells were then labeled with 5 µl Annexin V-fluorescein isothiocyanate (FITC) for 15 min at room temperature,5 µl propidium iodide (PI) for 5 min after being suspended in 300µl 1X binding buffer in the dark.Subsequently, apoptosis was detected using a flow cytometer (Beckman; Beckman Coulter, Inc., USA) after added 200µl 1X binding buffer.And the Annexin V and PI values were set as the horizontal and vertical axes, respectively, for the plot construction.Mechanically damaged, late apoptotic, dual negative/normal, and early apoptotic cells were located in the upper left, upper right, lower left and lower right quadrants of the flow cytometric dot plot, respectively.

#### Serum and mouse renal epithelial cells supernatant TNF-α, IL-1β and SCr levels measurement by Enzyme linked immunosorbent assay (ELISA)

The upper serum of mice in each animal experiment groups, the cell supernatant of each mouse renal epithelial cell model was taken respectively, and the both were tested separately.The levels of cytokines including TNF-α,IL-1β and SCr were measured with commercially available ELISA kits (Mlbio, Shanghai, China), according to the manufacturer’s instructions. The concentrations of cytokines were calculated by the standard curve.

#### Measurement of Bax and Bcl-2 in mouse renal epithelial cells

Cells in each group were collected, then total proteins were extracted and the concentration was determined by using Pierce™ BCA Protein Assay Kit. Proteins were separated on 10% sodium dodecyl sulphate-polyacrylamide gel electrophoresis (SDS-PAGE) gel and transferred onto polyvinylidene difluoride (PVDF) membranes.Then, the membranes were blocked with 5% nonfat milk in the Tris-buffered saline-Tween (TBST) solution for 1 hour and incubated overnight at 4°C with anti-rat polyclonal primary antibodies to Bax (1:5000, Proteintech, Chicago, USA), Bcl-2 (1:1000, Proteintech, Chicago, USA),respectively. After washed with TBST for 3 times, the membranes were incubated with goat anti-mouse secondary antibodies for conjugating with horseradish peroxidase (HRP) for 2 hours.Specific bands were visualized by using ECL chemiluminescent detection system and the expression levels were quantized by Bandscan software.

#### Statistical Analysis

All data was presented as mean ± SD. The statistical analysis was performed with SPSS 25.0 (SPSS Inc. IBM, Armonk, NY, USA). The results were evaluated in the Kolmogorov-Smirnov (KS) test of normality and analyzed with one-way ANOVA Student-Newman-Keuls test in different groups. Significant differences were considered when p-values < 0.05.

## Results

### The characteristics of exosomes

After extracted, the exosomes were observed with an electron microscope, and the morphology was shown in Figure 1.The particle size analyzed with DLS was concentrated at 95nm.

**Figure 1:**
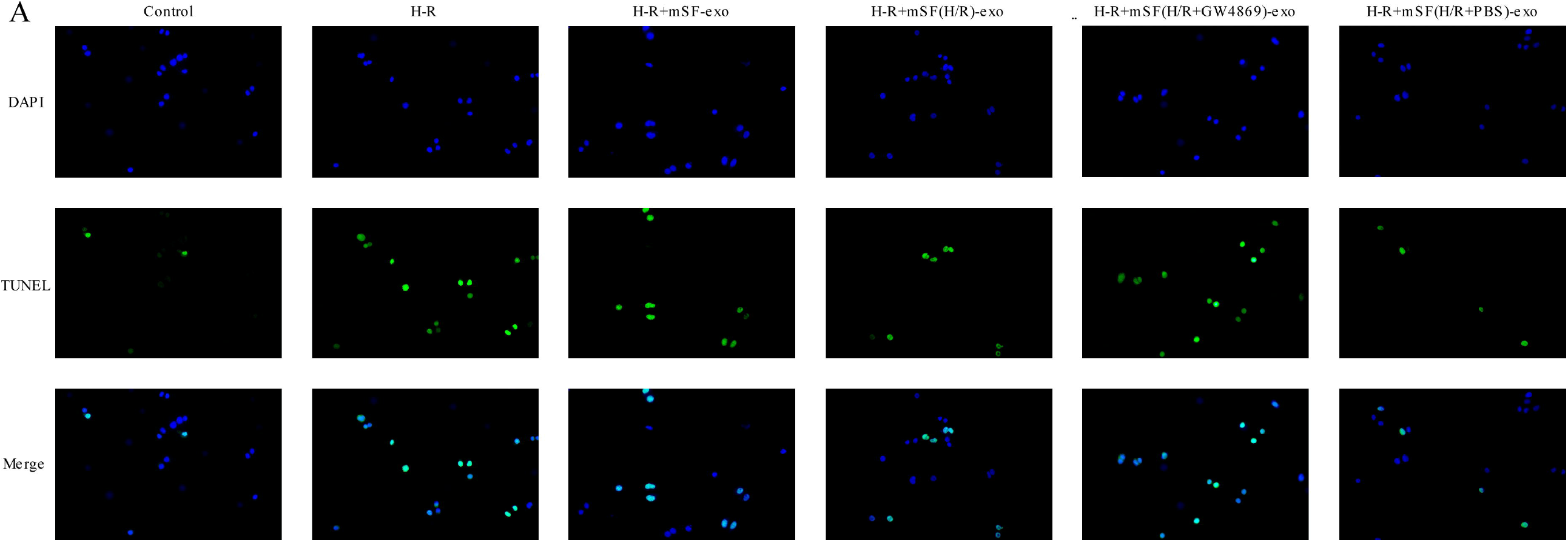
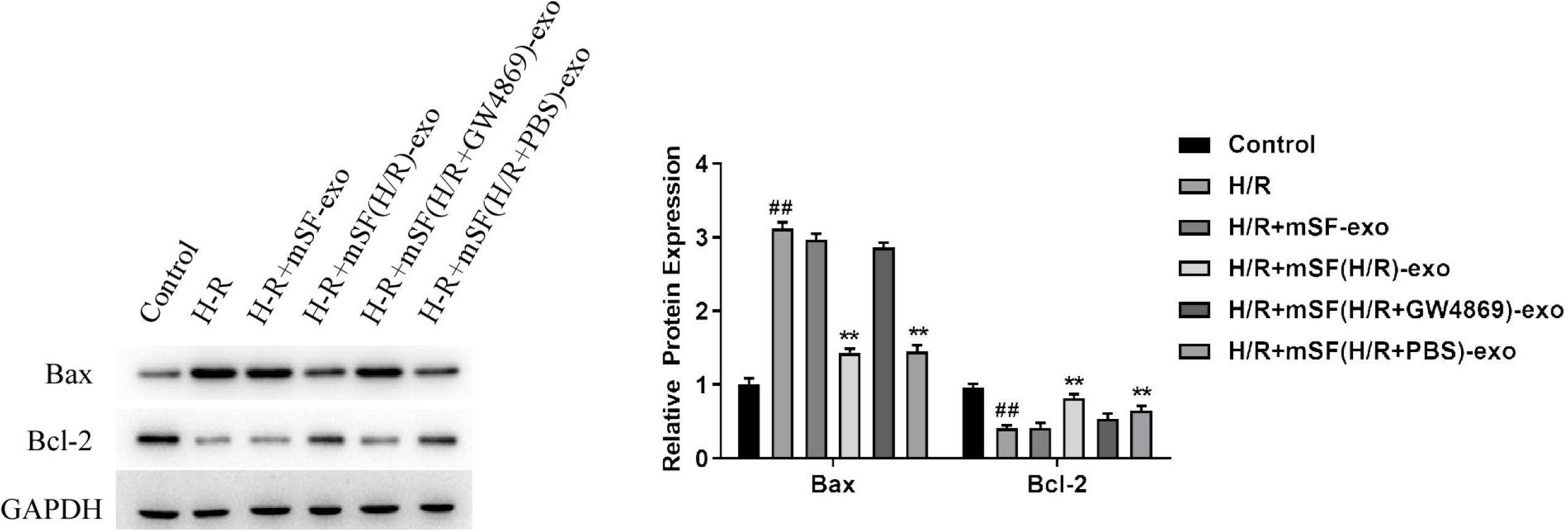
The characteristics of exosomes.(A) The morphology of exosomes under the electron microscope;(B)The particle size of exosomes analyzed with DLS.

### Exosomes extracted from mSF H/R cell model relieve renal ischemia-reperfusion injury

For all mouse kidney tissue sections, we performed hematoxylin-eosin(HE)staining, ki67 immunohistochemistry and TUNEL staining.

We found that there are differences in inflammation between different groups in histologic analysis by HE staining(Figure 2A). The cells in the sham group arrange intactly and are in normal morphology. The renal tissues in the IRI group are hyperemic, with deep stained nuclei and a large amount of inflammatory cells infiltration. The pathological changes in the IRI+mSF(H/R+GW4869)-exo group are similar to those in the IRI group. IRI+mSF-exo group has milder lesions than IRI group, with moderate inflammatory cells infiltration. The pathological changes in the IRI+sIPC group, IRI+mSF(H/R)-exo group, IRI+mSF(H/R+PBS)-exo group are similar with mild inflammatory cells infiltration, lighter than the IRI group, more serious than the sham group.

**Figure 2:**
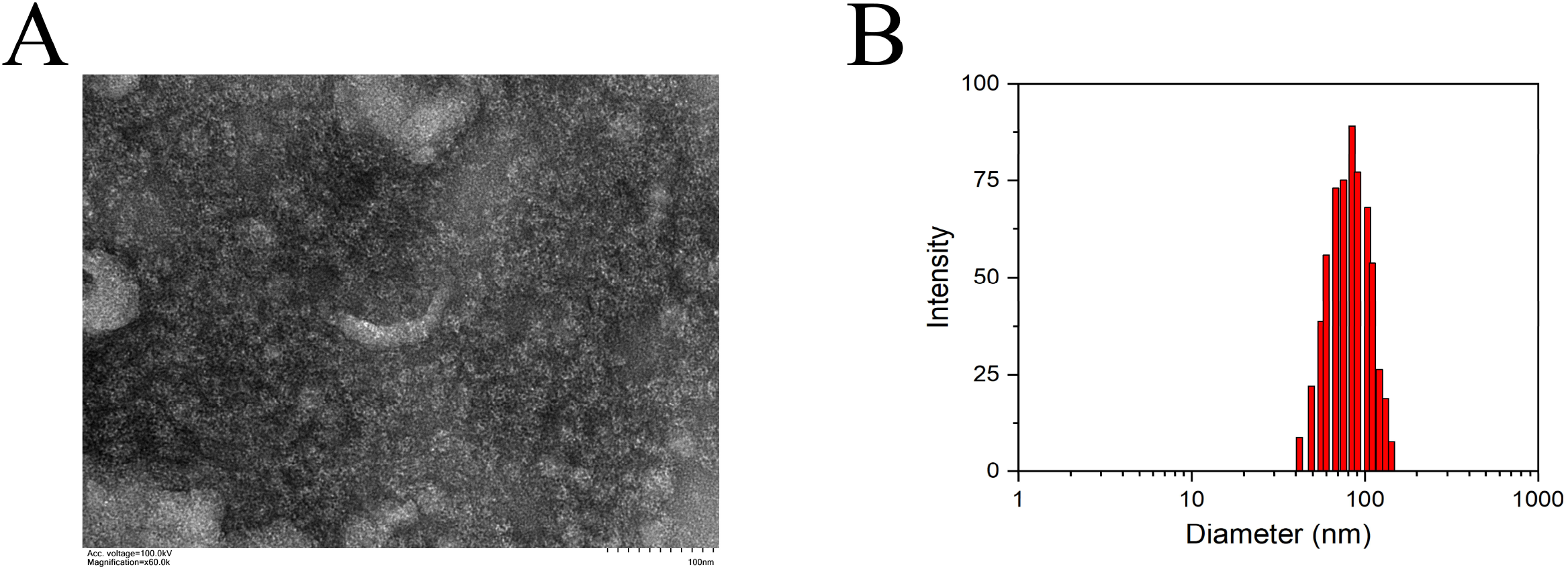
Microscopic images of the kidney tissues. (A)Hematoxylin-eosin(HE) staining;(B)Ki-67 immunohistochemistry;(C)TUNEL test;Pathological examinations showed that renal ischemia-reperfusion caused obvious renal injury in mice and there were more TUNEL staining positive cells and less Ki67 expression cells, indicating that renal ischemia-reperfusion caused increased cell damage and decreased cell proliferation. The exosomes which were extracted from splenic ischemic preconditioning can alleviate the injury caused by renal ischemia-reperfusion while the use of exosome release inhibitor inhibited the protective effects.

Immunohistochemical analysis was used to evaluate the levels of protein expression of Ki-67 which revealed cell proliferation in the kidney(Figure 2B).The expression of Ki-67 protein in the IRI group and IRI + mSF(H/R+GW4869)-exo group are similar and the weakest. The expression in the sham group is the strongest, and then the expression in the IRI+mSF-exo group is only weaker than the IRI group. The expression in the IRI+sIPC group, IRI+mSF(H/R)-exo group, IRI+mSF(H/R+PBS)-exo group are about the same, between the IRI and the sham group.

As shown in the TUNEL test(Figure 2C),the apoptotic cells in the sham group were rarely observed, but a very large number of apoptotic cells were found in the IRI group which are close to the IRI+mSF(H/R+GW4869)-exo group;Compared with the IRI group, the apoptotic cells were much fewer after sIPC treatment; while less slightly after the tail vein injection of exosomes;the apoptotic cells in the IRI+mSF-exo group was lower than IRI group but higher than IRI+sIPC group.The number of the apoptotic cells in the IRI+sIPC group, IRI+mSF(H /R)-exo group, IRI+mSF(H/R+PBS)-exo group were close, which was lower than the IRI group but higher than the sham group.

### Inhibition of exosomes extracted from splenic ischemic preconditioning on TNF-α, IL-1β and SCr release and expression

The inflammatory response and renal function after renal I/R were determined by measuring the protein concentrations of TNF-α,IL-1β and SCr with ELISA in the kidney tissue and mouse renal epithelial cell model.The cytokines as an indicative of inflammation in IRI group, IRI+mSF-exo group and IRI + mSF(H/R+GW4869)-exo group were markedly increased compared to sham group (p<0.05)(Figure 3A).While, there was a significant decrease of them in IRI+sIPC group, IRI+mSF(H/R)-exo group, IRI+mSF(H/R+PBS)-exo group in comparison with IRI group (p<0.05),but still higher than sham group.In consistent with the serological results in mice, the levels of TNF-α and IL-1β in cell model were up-regulated in H/R group (p<0.05) and were down-regulated in H/R+mSF(H/R)-exo group, H/R+mSF(H/R+PBS)-exo group (p<0.05),which suggested that the releases and expressions of TNF-α and IL-1β were inhibited (Figure 3B).

**Figure 3:**
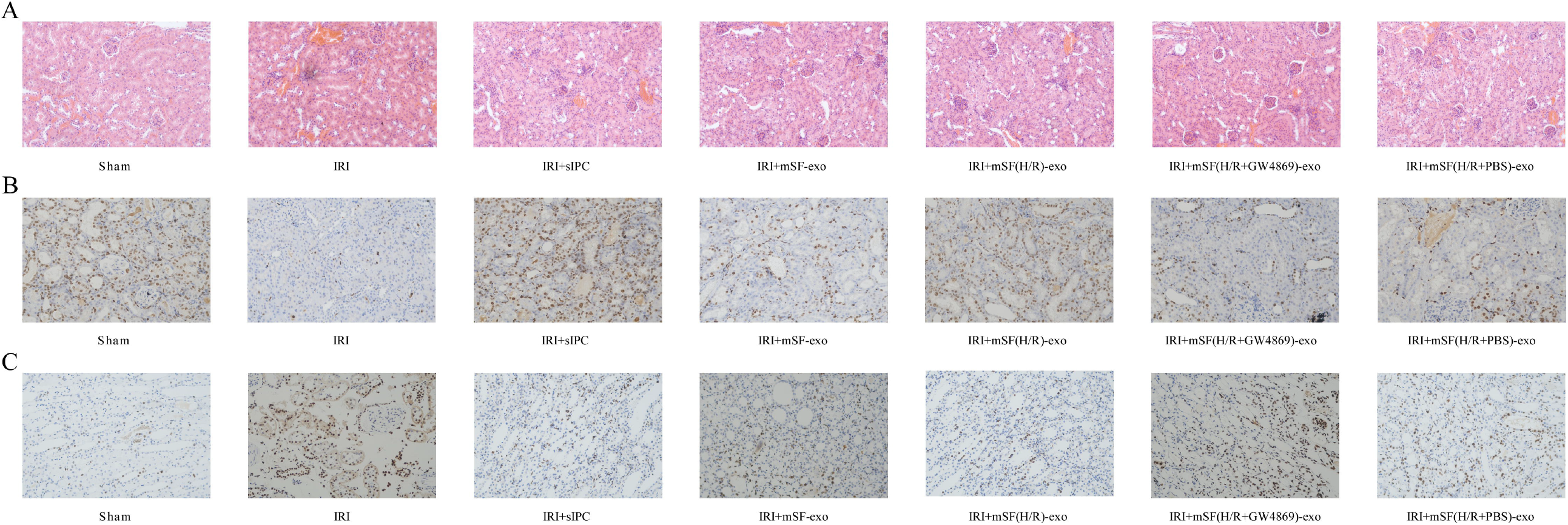
Effects of exosomes extracted from splenic ischemic preconditioning on inflammatory responses and renal function in mice and cellular models.(A)Differences in TNF-α, IL-1β and SCr levels between mouse model groups.(B)Differences in TNF-α,IL-1β between cell model groups.The values are presented in Mean ± SD; statistical analysis was carried out by one-way ANOVA Student-Newman-Keuls test,** means p<0.01,*** means p<0.001.

### Exosomes extracted from splenic ischemic preconditioning inhibit the apoptosis induced by IRI in mouse renal epithelial cell model

To confirm the anti-apoptotic effect of exosomes extracted from splenic ischemic preconditioning, flow cytometry, DAPI and TUNEL staining were performed. As shown in figure 4A,the apoptotic cells in control group was the lowest, H-R group and H-R+mSF(H/R+GW4869)-exo group were close to and the highest, and then H-R+mSF-exo group was the second highest.The apoptotic cells of H-R+mSF(H/R)-exo group and H-R+mSF(H/R+PBS)-exo group was the second lowest and only higher than control group.Furthermore, using flow cytometry analysis, we verified the same result (figure 4B).

**Figure 4:**
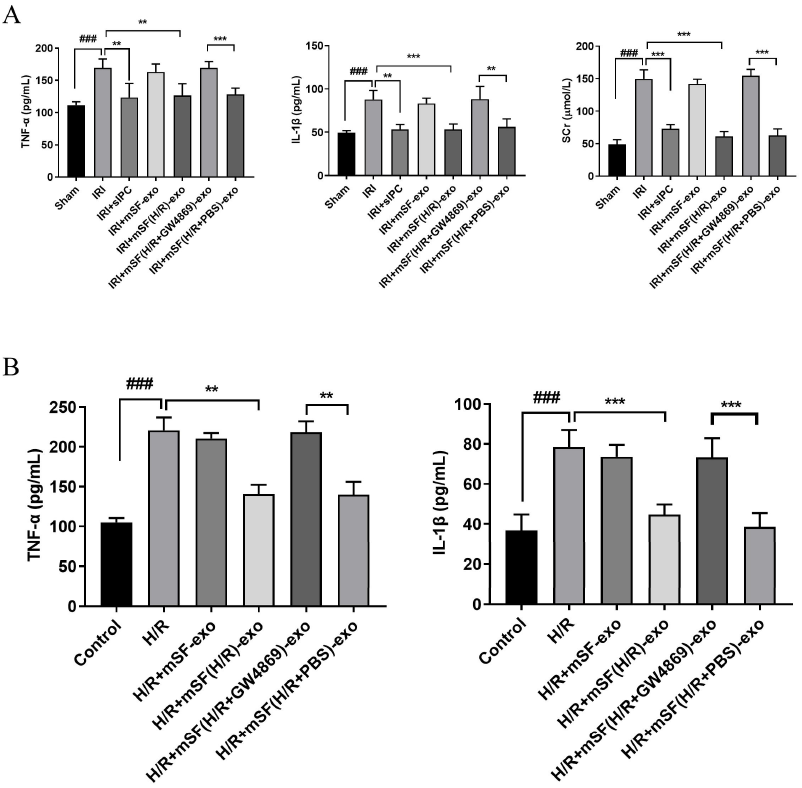
The exosomes have a protective effect on renal epithelial cell apoptosis induced by ischemia-reperfusion.(A)For DAPI and TUNEL staining, the number of TUNEL-positive cells in H/R group was higher.After stimulation of exosomes extracted from starvation-treated mSF, the TUNEL-positive cells decreased, while elevated significantly after the addition of exosome release inhibitor.(B)Flow cytometry showed that there were more apoptotic cells in H/R group, and starvation-treated splenocyte exosomes reduced the injury induced by ischemia-reperfusion.

### Exosomes extracted from splenic ischemic preconditioning up-regulated the anti-apoptotic protein Bcl-2 and down-regulated the pro-apoptotic protein Bax

Western blot revealed that the expression levels of Bcl-2 were decreased while Bax were increased significantly in H-R group. Obvious that, Bax was the lowest in control group, and the highest in H-R group. The expression of Bax in H-R +mSF-exo group and H-R +mSF(H/R+GW4869)-exo group was close and the second highest.However, the Bax in H-R +mSF(H/R)-exo group and H-R +mSF(H/R+PBS)-exo group was close and the second lowest.The difference of Bcl-2 in each group is completely opposite to that of Bax(Figure 5).

**Figure 5:**
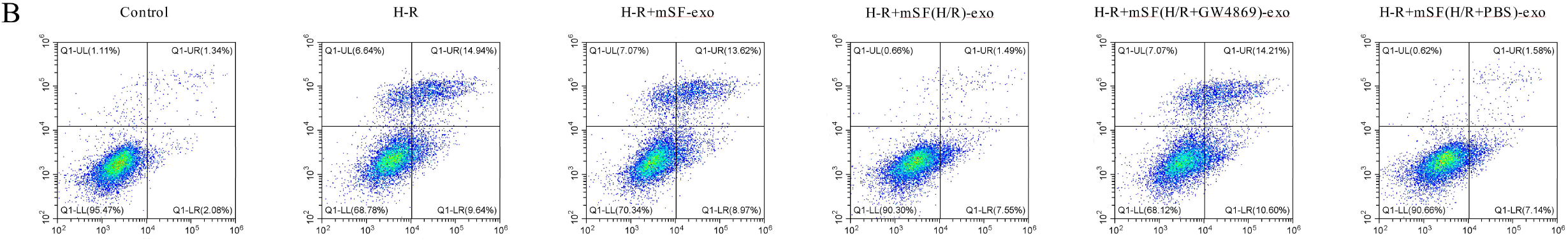
Western blot analysis of Bcl-2 and Bax in each cell model group. **,## means p<0.01.

## Discussion

Ischemia reperfusion injury is common in transplantation^[11]^, partial nephrectomy^[12, 13]^ an shock. The short-term ischemic preconditioning of an organ can make the distant organs adapt to changes in advance, reduce the pressure of ischemia-reperfusion on blood vessels and target organs, and then prevent the damage caused by the subsequent long-term ischemia-reperfusion.Renal ischemia-reperfusion injury has a great influence on its function preservation. Our previous studies have confirmed that splenic ischemic preconditioning can protect the kidney from IR injury by inhibiting the NF-κB pathway and reducing the expression of pro-inflammatory cytokines^[6]^.

This study further explored the specific role of exosomes in this inhibition. As a naturally equipped biological carrier, exosomes play an important role not only between intercellular but also inter-organ communication^[14, 15]^. More and more studies have shown that exosomes are of great significance for reducing the ischemia-reperfusion injury of the heart^[16, 17]^, brain^[18]^, spinal cord^[19]^, liver^[20]^ and other organs. Li X et al.^[21]^ found miR-146a-5p which was the the most abundant miRNA in exosomes targeted and degraded the 3’UTR of interleukin-1 receptor-associated kinase 1 (IRAK1) mRNA, and then inhibited nuclear factor (NF)-κB signaling and inflammatory cell infiltration to reduce renal ischemia-reperfusion injury. Kim S et al.^[22]^used optogenetically engineered exosome technology deliver the exosomal super-repressor inhibitor of NF-ĸB (Exo-srIĸB) into mice before/after kidney ischemia-reperfusion surgery.Subsequently the gene expression of pro-inflammatory cytokines and adhesion molecules decreased, proving that exosomes were effective in limiting renal ischemia-reperfusion injury.

These facts give us a hint that exosomes may act as an information medium between the spleen and kidney and then alleviate renal ischemia-reperfusion injury.We hypothesized that exosomes derived from the spleen can act as a medium for information dissemination in a certain way, further transfer to the kidney tissue. In order to prove our inference, we purified the exosomes from the hypoxia-reoxygenation model of splenic fibroblasts and infused them intravenously into mice with renal ischemia-reperfusion injury.In vitro, the exosomes were applied on the hypoxia-reoxygenation model of renal epithelial cells. Furthermore, HE staining, immunohistochemistry, TUNEL staining, ELISA detection, flow cytometry, western blot and other techniques were used to compare the differences between various groups. The results showed that the exosomes extracted by splenic ischemic preconditioning can effectively induce cell apoptosis inhibition and reduce renal injury and oxidative stress, while the addition of exosomes release inhibitors weakens the protective effects in the I/R model.

As in our previous study^[6]^, compared with sham group, the sCr level of mice after ischemia-reperfusion was significantly increased, suggesting that IRI leads to impaired renal function. In IRI+sIPC group, the kidney underwent ischemia-reperfusion experiments after splenic ischemic preconditioning in which the spleen underwent three cycles of ischemia for 5 minutes and reperfusion for 5 minutes. The sCr significantly decreased, which means that splenic ischemic preconditioning has a protective effect on renal IR injury. In the animal experiments of this project, we evaluated the effect of exosomes extracted from splenic ischemia preconditioning on renal IR injury by HE staining, ki67 immunohistochemistry, and TUNEL staining. Finally, we confirmed the protective effect of exosomes.

Following ischemia-reperfusion injury, neutrophil infiltration increases and pro-inflammatory cytokines such as TNF-α and IL-1β are activated^[23]^.The increase of pro-inflammatory cytokines will promote the phosphorylation of NF-κB, and its activation/phosphorylation can enhance the pro-inflammatory response and further lead to the renal injury^[24]^. Therefore, we evaluated the levels of TNF-α and IL-1β, both of which significantly reduced after splenic ischemic preconditioning or after the addition of the exosomes extracted from mSF(H/R). This inhibitory effect on the pro-inflammatory cytokines demonstrates that exosomes extracted from splenic ischemic preconditioning play the protective effect by their anti-inflammatory properties.

As the key regulators of pathways involved in cell apoptosis, Bcl-2 family proteins play a role in inhibiting or promoting cell death, among which Bcl-2 and Bcl-XL are anti-apoptotic factors, while other members such as Bax and Bcl-Xs promote apoptosis^[25]^.The increase of Bax and the decrease of Bcl-2 will destroy the integrity of mitochondria, leading to the release of cytochrome C^[26, 27]^.And then apoptotic bodies form with caspase-9,activating the downstream caspase cascade, inducing apoptosis of renal tubular cells and leading to impaired renal function. We found that starvation-treated spleen fibroblasts exosomes can reduce Bax and increase Bcl-2, the findings will provide a new view and evidence for the protective effect of exosomes in the case of renal I/R injury.

In conclusion, we confirmed that the exosomes extracted after splenic ischemic preconditioning had a protective effect on renal ischemia-reperfusion injury from two aspects of animal model and cell model.

## Conclusions

We have confirmed that exosomes extracted from splenic ischemic preconditioning have a protective effect on renal IR injury. Potentially, exosomes extracted from splenic ischemic preconditioning can serve as a promising therapeutic option for renal IR.However, there were some limitations in this study. It wasn’t mentioned which components in the exosomes play the protective role and the specific signaling pathways. These problems need further research and discussion.

